# Abnormal function in dentate nuclei precedes the onset of psychosis: a resting-state fMRI study in high-risk individuals

**DOI:** 10.1101/2021.02.28.433240

**Authors:** Sheeba Arnold Anteraper, Xavier Guell, Guusje Collin, Zhenghan Qi, Jingwen Ren, Atira Nair, Larry J. Seidman, Matcheri S. Keshavan, Tianhong Zhang, Yingying Tang, Huijun Li, Robert W. McCarley, Margaret A. Niznikiewicz, Martha E. Shenton, William S. Stone, Jijun Wang, Susan Whitfield-Gabrieli

## Abstract

**Objective:** The cerebellum serves a wide range of functions and is suggested to be composed of discrete regions dedicated to unique functions. We recently developed a new parcellation of the dentate nuclei (DN), the major output nuclei of the cerebellum, which optimally divides the structure into three functional territories that contribute uniquely to default-mode, motor-salience, and visual processing networks as indexed by resting-state functional connectivity (RsFc). Here we test for the first time whether RsFc differences in the DN precede the onset of psychosis in individuals at risk of developing schizophrenia.

**Methods:** We used the MRI dataset from the Shanghai At Risk for Psychosis study that included subjects at high risk to develop schizophrenia (N=144), with longitudinal follow-up to determine which subjects developed a psychotic episode within one year of their fMRI scan (converters N=23). Analysis used the three functional parcels (default-mode, salience-motor, and visual territory) from the DN as seed regions of interest for whole-brain RsFc analysis.

**Results:** RsFc analysis revealed abnormalities at baseline in high-risk individuals who developed psychosis, compared to high-risk individuals who did not develop psychosis. The nature of the observed abnormalities was found to be anatomically specific such that abnormal RsFc was localized predominantly in cerebral cortical networks that matched the three functional territories of the DN that were evaluated.

**Conclusions:** We show for the first time that abnormal RsFc of the DN may precede the onset of psychosis. This new evidence highlights the role of the cerebellum as a potential target for psychosis prediction and prevention.

## INTRODUCTION

A large and expanding body of evidence has demonstrated that cerebellar structural and functional abnormalities exist in patients diagnosed with neurological and psychiatric disorders that degrade cognition and affect, including Alzheimer’s disease (1), Parkinson’s disease (2), frontotemporal dementia (1), major depressive disorder (3, 4), autism spectrum disorder (ASD) (5, 6), attention-deficit/hyperactivity disorder (ADHD) (7), dyslexia (8) and schizophrenia (9, 10). Several of these conditions are neurodevelopmental in nature (e.g., ADHD, ASD, dyslexia) and show abnormalities early on. However, much less is known about whether these cerebellar abnormalities precede the manifestation of symptoms in neurodevelopmental conditions with later symptom onsets such as schizophrenia. Existing evidence of cerebellar abnormalities in patients diagnosed with neuropsychiatric disorders improves our understanding of their pathophysiology and may open new avenues for their diagnosis and treatment. The question of whether cerebellar disruption precedes the onset of symptoms in these conditions constitutes a fundamental knowledge gap in neuropsychiatry; addressing this gap will allow investigation of the cerebellum as a potential target for disease prediction and prevention.

Here we aim to test whether abnormalities in the dentate nuclei (DN), a collection of neural bodies buried beneath the cortex of each cerebellar hemisphere (**Figure 1**), precede the onset of symptoms in individuals at risk for schizophrenia. Anatomical connections between cerebellar cortex and extracerebellar territories engaged in thought and affect form the basis of the neuroscience of cerebellar behavioral neurology and psychiatry (11, 12). The DN are central structures in these anatomical pathways, as the majority of the fibers that exit the cerebellar cortex synapse in the DN before reaching extracerebellar structures such as thalamus or cerebral cortex (13). To the best of our knowledge, this is the first study assessing if DN abnormalities precede the onset of symptoms in individuals at risk for schizophrenia.

**Figure 1:**
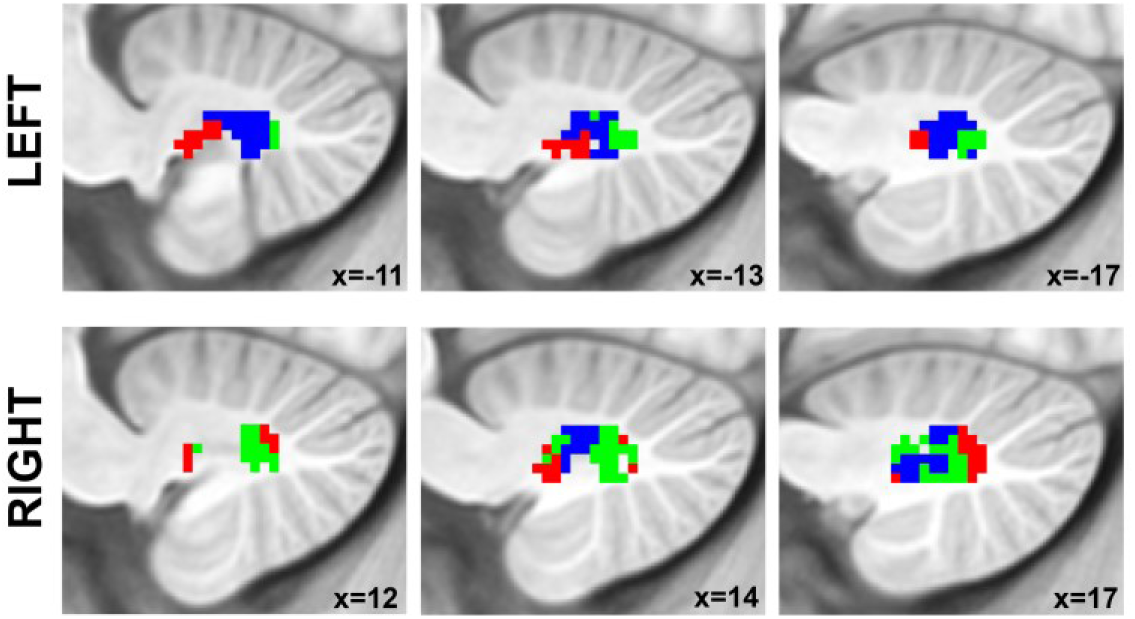
Structural location and functional territories of the dentate nuclei as reported in (43). Red = default-mode network functional territory. Blue = salience-motor functional territory. Green = visual functional territory.

The evidence for cerebellar abnormalities in schizophrenia is limited in comparison to other brain areas (14) because the cerebellum was long overlooked in psychiatric research due to erroneous notions that it is not involved in higher cognitive functions and cognitive-behavioral disorders (15). The past two decades, however, have seen a growing interest in the role of the cerebellum in schizophrenia, led by Nancy Andreasen’s work on the role of prefrontal-thalamic-cerebellar circuitry in schizophrenia and her cognitive dysmetria hypothesis (16, 17) following Schmahmann’s dysmetria of thought theory (12, 18). Neuroimaging studies since then have shown consistent evidence for cerebellar abnormalities in schizophrenia. Task-activation studies indicate both hypo- and (to a lesser extent) hyper-activation of the cerebellum in a range of tasks (19, 20), with review/meta-analytic data suggesting that hypoactivations predominate in medial portions of the anterior lobe and lobules IV and V, while hyperactivations localize more laterally in lobules VI and VII (20) and that abnormalities in task-activation may stem from an altered functional topography of the cerebellum (19). In addition, functional connectivity studies provide robust evidence for abnormalities in cortico-cerebellar and thalamo-cerebellar functional connectivity in patients with schizophrenia and high-risk individuals (10, 21–30). These studies also report both hypo- and hyperconnectivity for different cerebellar regions and distinct cerebro-cerebellar connections, supporting the diverse functional roles and heterogenous connectivity profiles of individual cerebellar regions (27). Across studies, there appears to be relatively consistent evidence for cerebro-cerebellar hyperconnectivity in some networks including the somatomotor and default-mode network (DMN) (23, 25, 26, 29), while findings for other functional networks are more mixed. For example, some studies report hypoconnectivity between regions of the ventral attention or salience network and corresponding cerebellar clusters (26, 29) while others report hyperconnectivity for similar cerebro-cerebellar connections (27). Recent studies have also shown evidence of abnormal cerebellar functional connectivity in schizophrenia as indexed by functional gradients analyses, highlighting the possibility of large-scale disruptions in communication within and between all major functional territories of the cerebellar hemispheres and their communication with the cerebral cortex (31).

Neuropathological studies have also found evidence for cerebellar pathology in schizophrenia (32), including findings of a decreased density (33) and size (34) of Purkinje cells. The latter finding is particularly significant as Purkinje cells provide input to deep nuclei including the DN, and thereby play a key role in modulating the output from the cerebellum to the cerebral cortex (17, 34). Moreover, studies in mice show that stimulation of the DN modulates prefrontal dopamine, a neurotransmitter system implicated in schizophrenia (35). This may be part of the mechanism underlying emerging findings that cerebellar stimulation improves symptoms of schizophrenia (36, 37). Taken together, existing evidence suggests that schizophrenia involves an altered functional modulation of higher cortical functions by the cerebellum. As the main output nucleus from cerebellum to the cerebral cortex, the DN may play a key role in this process and may thus be a promising target for intervention. These findings point to the importance of investigating dentate functional anatomy in schizophrenia, but there are no functional MRI studies to date that focus on the DN and its functional connectivity to the cerebral cortex in individuals diagnosed with or at risk for schizophrenia.

The natural history of schizophrenia makes it possible to test whether at-risk individuals reveal DN functional abnormalities that precede the formal onset of the disease. The dataset employed here included subjects at clinical high risk to develop psychosis, with longitudinal follow-up to determine which subjects developed a psychotic episode within one year of their fMRI scan, as described in (38) and determined by standardized diagnostic scales in psychiatry. The prodromal stage is considered a period of imminent risk for psychosis. The clinical high risk paradigm was developed based on observations that the majority of patients with schizophrenia experience a prodromal phase characterized by attenuated or transient psychotic symptoms including unusual thought content, suspiciousness, or mild perceptual abnormalities in the months to years preceding the first psychotic episode. Around 20% of individuals showing such subthreshold symptoms convert to psychosis within 2 years (39). To compare, the annual incidence rates of psychosis in the general population are estimated around 0.5 per 1000. Conversion rates in our current sample are similar to those reported in literature (39, 40), with approximately 16% developing a psychotic episode by one-year follow-up. Of note, our current sample is a subset of the total study sample for whom good-quality fMRI data was available. Clinical profiles and conversion rates on the total sample have been described in detail previously (41). There is a relatively paucity of studies on outcomes in high-risk non-converters, but it is thought that approximately half maintain stable levels of subthreshold symptoms, while the other half show remission of subthreshold symptoms (42).

Contrasting prior investigations using the whole DN as one single region of analysis, our group recently showed that human DN are optimally divided into three functional territories that contribute uniquely to DMN, motor-salience, and visual processing as indexed by resting-state functional connectivity (RsFc) MRI (43) (**Figure 1**). This improved understanding of a crucial node of cerebellar functional anatomy that can be captured with RsFc, combined with longitudinal data differentiating high-risk subjects who develop psychosis versus high-risk subjects who do not develop psychosis, provides a novel avenue to study functional abnormalities in DN at the earlier stage of pathophysiological development.

## METHODS

### Study Participants

The study participants were individuals at Clinical High Risk (CHR) for psychosis (N=144; mean age = 19, age range=13–34) from the Shanghai At Risk for Psychosis (SHARP) program, an international collaborative research effort to recruit a unique sample of medication-naïve adolescents and young adults with early signs of impending psychosis. Twenty-three of the CHR subjects developed psychosis before 1-year clinical follow-up (CHR+). Ninety-three age-, gender-, education-, and handedness-matched healthy controls (HC) (mean age = 18.7, age range=12–35) were also part of this dataset. Clinical characteristics, conversion criteria, and demographics information have been previously reported (38). Briefly, a Chinese version of the Structured Interview for Prodromal Symptoms (SIPS (44)) was used to assess prodromal symptoms. There were no significant differences in baseline SIPS scores in CHR+ compared to CHR participants who did not develop psychosis (CHR-). Institutional Review Boards at Beth Israel Deaconess Medical Center and the Shanghai Mental Health Center approved the study. Informed consent was obtained from all participants or their legal guardians. DSM-IV Alcohol or Drug Dependence within 3 months of study participation was an exclusion criterion to our study. IQ< 70, and any medical condition, sensorimotor handicap, or acquired injury that may either contribute to prodromal symptoms or confound prodromal symptom assessment (e.g., deadness, blindness, acquired brain injury, epilepsy) were also grounds for exclusion. Detailed demographic and clinical characteristics are provided in **Table 1**.

**Table 1.** Demographic and clinical characteristics

|  | CHR+<br>(*N* = 23) | CHR-<br>(*N* = 121)<sup>a</sup> | HC<br>(*N* = 93) | Statistics |
| --- | --- | --- | --- | --- |
| Age in years, mean (sd)<br>[range] | 19.2 (5.2)<br>[14 – 34] | 18.8 (5.0)<br>[13 – 32] | 18.7 (4.6)<br>[12 – 35] | $F = 0.10, p = 0.91$ |
| Sex, male/female | 16 / 7 | 57 / 64 | 49 / 44 | $\chi = 3.99, p = 0.14$ |
| Education in years,<br>mean (sd) [range] | 10.3 (2.2)<br>[7 – 16] | 10.6 (3.0)<br>[4 – 19] | 10.8 (2.3)<br>[6 – 17] | $F = 0.35, p = 0.71$ |
| IQb, mean (sd) [range] | 92.1 (19.0)<br>[52 – 112] | 100.1 (11.7)<br>[67 – 128] | 104.2 (11.1)<br>[75 – 133] | $F = 8.06, p < 0.001$ |
| *Baseline SIPS scores* |  |  |  |  |
| Positive, mean (sd)<br>[range] | 10.0 (3.3)<br>[4 – 17] | 10.1 (3.8)<br>[0 – 21] | | $F = 0.03, p = 0.87$ |
| Negative, mean (sd)<br>[range] | 12.1 (6.4)<br>[3 – 26] | 11.5 (6.0)<br>[1 – 27] | | $F = 0.24, p = 0.63$ |
| Total, mean (sd) [range] | 37.6 (10.7)<br>[16 – 65] | 37.2 (11.0)<br>[13 – 79] | | $F = 0.02, p = 0.90$ |
| *Psychotropic medication* |  |  |  |  |
| At inclusion, *N* (%) | 1 (4.3%) | 6 (5.0%) | | $\chi = 0.12, p = 0.90$ |
| At baseline MRI, *N* (%) | 7 (30.4%)<sup>c</sup> | 21 (17.4%) | | $\chi = 2.11, p = 0.15$ |
| Antipsychotics, *N* (%) | 6 (26.1%) | 17 (14.0%) | | $\chi = 2.09, p = 0.15$ |
| Antidepressants, *N* (%) | 2 (8.7%) | 4 (3.3%) | | $\chi = 1.41, p = 0.24$ |
| Anxiolytics, *N* (%) | 1 (4.3%) | 0 (0%) | | $\chi = 5.30, p = 0.02$ |

<sup>a</sup> Of note, the CHR- group is smaller than in a previous publication on this sample (Collin et al., 2018), because the current study excluded 14 CHR- who were lost to follow-up. <sup>b</sup> IQ data were unavailable for 4 CHR+, 16 CHR-, and 9 HC. <sup>c</sup> The number of CHR converters on psychotropic medication at the time of baseline MRI (*N* = 7) is lower than the sum across medication types, because two individuals were on two different medications (i.e., combining an antipsychotic with an antidepressant or anxiolytic).

### Structural and Resting-state Functional MRI Acquisition Parameters

Imaging data were collected on Siemens 3T MR B17 (Verio) system, with Siemens 32-channel head coil. High-resolution whole-brain structural data (1 mm isotropic voxels) were acquired using a sagittal T1-weighted MPRAGE sequence with duration 9 min 14 s. Scan parameters for TR/TE/Flip Angle were 2.3 s/2.96 ms/9°. Whole-brain resting-state functional data (3.5 mm isotropic voxels) were acquired using a T2*-weighted EPI sequence with duration 6 min 19 s. Scan parameters for TR/TE/Flip Angle were 2.5 s/30 ms/90°, 37 contiguous slices.

### Data processing: Seed-to-voxel Functional Connectivity Analysis

Resting-state fMRI data were realigned and spatially normalized to the MNI template using SPM12 (Wellcome Department of Imaging Neuroscience; www.fil.ion.ucl.ac.uk/spm). Structural images were segmented into white matter (WM), gray matter, and cerebrospinal fluid (CSF) using SPM12. The CONN Toolbox (45) was used to compute whole-brain r-maps from the seed regions of interest (ROIs). ROIs, defined at the group level, included the whole DN (as defined using the SUIT DN mask; (46)), and three functional sub-territories of DN that were defined in a previous study by our group (43), including DMN, motor-salience, and visual functional regions (**Figure 1**). The CONN Toolbox uses an anatomical component-based correction method (aCompCor, (47)) for denoising BOLD time-series and integrates quality assurance methods (Artifact Detection Tools, www.nitrc.org/projects/artifact_detect). Band-pass filtering was carried out at 0.008 – 0.09 Hz. Time points with mean signal intensity outside three standard deviations from global mean signal, and 0.4 mm scan-to-scan motion (about 1/10^th^ the acquisition voxel size) were flagged as problematic scans. There was no significant between-group difference (CHR+ versus CHR-) in the number of time-points that were flagged as motion outliers (p=0.87) and these time-points were regressed out along with six realignment parameters and physiological sources of noise (three principal components of WM, and three principal components of CSF segments, using aCompCor; (47)). WM and CSF segments were derived from the structural images using the segmentation routine in SPM12. Because of the small size of the DN, unsmoothed data was used for data analysis to minimize partial volume effects from structures close to DN. This strategy has been employed previously for functional connectivity analysis of dentate ROIs (48). Whole brain Pearson’s correlation maps derived from denoised time-series from whole DN and the three DN functional territories were converted to z-scores using Fischer’s r to z transformation to carry out second-level general liner model (GLM) analyses.

### Data processing: Second Level GLM Analysis

For all three groups (HC, CHR+, and CHR-), seed-to-voxel analysis was carried out using the whole DN as a seed, as well as using the unique effect of each of the three functional territories (DMN, salience-motor, and visual). The unique effect of each functional territory was calculated using the same analysis method as in (43), i.e., the DMN unique effect was calculated as DMN>(salience-motor and visual), salience-motor unique effect was calculated as salience-motor>(DMN and visual), and visual unique effect was calculated as visual>(DMN and salience-motor). Statistical significance thresholding for three-group effects (using ANOVA) included p < 0.005 (two-sided) at the voxel level and p < 0.05 False Discovery Rate (FDR) correction at the cluster level.

## RESULTS

### Second Level GLM Analysis

When using the whole DN as a seed, group contrasts between CHR+ and CHR-did not show statistically significant differences. In contrast, statistically significant differences were detected for each of the three functional sub-regions in DN (DMN, salience-motor, and visual). For the DN-DMN functional territory, the CHR+ group showed increased functional connectivity with the posterior cingulate cortex (PCC), right angular gyrus (AG), and dorsolateral prefrontal cortex (DLPFC); and decreased functional connectivity with primary and supplementary motor areas (**Figure 2A**). For the DN salience-motor functional territory, the CHR+ group showed increased functional connectivity with the postcentral gyrus; and decreased functional connectivity with the angular gyrus (**Figure 2B**). For the DN-visual functional territory, the CHR+ group showed increased functional connectivity with the dorsal anterior cingulate cortex; and decreased functional connectivity with precuneus / visual association areas, as well as DLPFC (**Figure 2C**). Cluster statistics are reported in **Table 2**.

**Figure 2.**
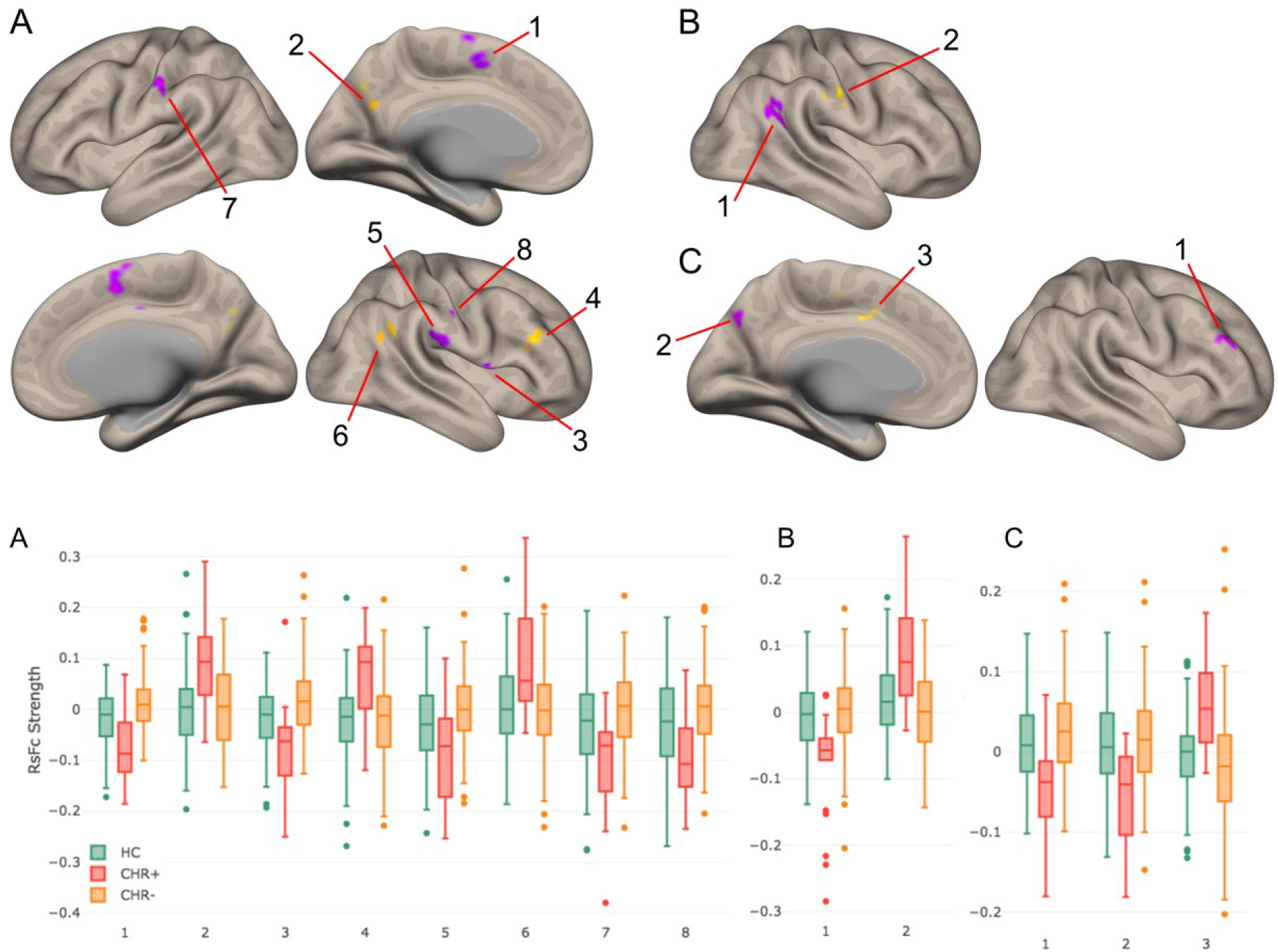
Top: Group contrast results (CHR+ versus CHR-) using functional connectivity calculated from the DMN functional territory of the DN (A), salience-motor functional territory of the DN (B), and visual functional territory of the DN (C), at voxel-level height threshold of p < 0.005 (two-sided) and cluster size FDR correction of p < 0.05. Bar plots at the bottom of the figure provide data for each cluster in CHR+ and CHR-, as well as data for the same clusters in the control group. Notably, for each of these group contrasts, data from CHR-participants were similar to healthy controls.

**Table 2.** Results from second-level seed-to-voxel analysis for CHR+ versus CHR-contrast for each functional territory within the DN (height-threshold = p<0.005 (two-sided); cluster threshold = p<0.05 FDR corrected).

| **Contrast:**<br>**CHR+ vs. CHR-** | **Region** | **Cluster**<br>**Size** | **Maximum**<br>**T-value** | **MNI**<br>**Coordinates (X,**<br>**Y, Z)** |  |  |
| --- | --- | --- | --- | --- | --- | --- |
| **DMN** | Supplementary motor cortex | 192 | -7.15 | 8 | 4 | 52 |
|  | Precuneus cortex / posterior cingulate cortex | 102 | 4.83 | 0 | -70 | 38 |
|  | Central opercular cortex | 75 | -6.06 | 52 | 6 | 0 |
|  | Dorsolateral prefrontal cortex (dlPFC) | 60 | 5.17 | 44 | 30 | 24 |
|  | Primary somatosensory cortex | 59 | -5.44 | 58 | -22 | 20 |
|  | Angular gyrus | 57 | 4.85 | 54 | -50 | 24 |
|  | Premotor cortex | 54 | -5.32 | -54 | -24 | 38 |
|  | Secondary somatosensory cortex | 46 | -5.20 | 58 | -20 | 38 |
| **Salience-Motor** | Angular gyrus | 116 | -5.92 | 48 | -50 | 22 |
|  | Postcentral Gyrus | 60 | 5.51 | 58 | -18 | 38 |
| **Visual** | dlPFC | 66 | -5.60 | 44 | 40 | 28 |
|  | Precuneus cortex / visual association areas | 63 | -5.66 | -2 | -74 | 36 |
|  | Dorsal anterior cingulate cortex | 47 | 5.54 | -2 | -2 | 42 |

Taken together, the anatomical specificity of the observed group-differences can be summarized as follows. For DMN and salience-motor territories of the DN, we observed hyperconnectivity to cerebral cortical areas belonging to the same functional network, and hypoconnectivity to cerebral cortical areas belonging to a different functional network, with the exception of DLPFC that was hyperconnected to the non-matching DMN territory of the DN. The opposite was true for the visual territory of the DN: we observed hypoconnectivity to cerebral cortical areas belonging to the same functional network, and hyperconnectivity to cerebral cortical areas belonging to a different functional network, with the exception of DLPFC that was hypoconnected to the visual functional territory of the DN.

## DISCUSSION

Here we show for the first time that abnormalities in functional connectivity between the DN and cerebral cortical areas may precede the onset of psychosis. This conclusion is supported by DN resting-state fMRI connectivity measurements in individuals with early signs of impending psychosis. Our data revealed abnormalities in those high-risk individuals who subsequently developed psychosis, compared to high-risk individuals who did not develop psychosis. Furthermore, our results reveal anatomical specificity in the distribution of these functional abnormalities, as abnormal functional connectivity was observed to be localized predominantly in cerebral cortical networks associated with the three functional territories of the DN that were evaluated. Evaluation of functional connectivity from whole DN as a single region of interest did not reveal significant between group differences. Group-differences were only detected when analyzing functional connectivity from each of the three functional sub-territories of the DN defined in the atlas developed previously by our group (43). DN functional sub-divisions are thus useful to detect DN functional differences in disorders of thought and affect. Taken together, this new evidence highlights the role of the DN as a potential target for disease prediction and prevention in neuropsychiatric disorders.

### Anatomical specificity of DN functional connectivity differences and possible pathophysiological interpretations

Our interpretation for our findings of the distinctive topography of the DN connectivity pattern between at-risk subjects who developed psychosis versus at-risk subjects who did not develop psychosis (**Figure 2**) is as follows. For DMN and salience-motor DN territories, two shared principles (a, b) and one exception (c) are observed. The first principle (a) is that hyperconnectivity was observed in cerebral cortical areas belonging to the same DN functional territory that was being examined. The DMN functional territory of the DN was hyperconnected to cerebral cortical areas associated with the DMN including posterior cingulate cortex and angular gyrus. Similarly, the salience-motor functional territory of the DN was hyperconnected to primary motor and motor association areas of the cerebral cortex. The second principle (b) is that hypoconnectivity was observed in cerebral cortical areas that did not belong to the same network as the specific DN functional territory that was being examined. DMN functional territory of the DN was hypoconnected to primary motor, motor association, and supramarginal gyrus, which belong to motor and task-positive networks, not DMN. Again, similarly, the salience-motor functional territory of the DN was hypoconnected to the angular gyrus, which belongs to DMN. The only exception (c) is that right DLPFC, which belongs to task-positive networks rather than DMN, was hyperconnected to the DMN territory of the DN. For the visual territory of the DN, we observed a reversal of these observations (a, b, c). Hypoconnectivity was observed with visual association areas; this finding is a reversal of observation (a), and hyperconnectivity was observed with the anterior cingulate cortex that is not linked to visual networks; this finding is a reversal of observation (b). Hypoconnectivity rather than hyperconnectivity was observed with DLFPC; since DLPFC does not belong to the visual network, this finding is a reversal of observation (c).

The pathophysiological significance of these observations is difficult to establish, as the group-differences observed here may represent pathological abnormalities, compensatory reorganizations, or a combination of both. The fact that DLPFC showed a reversed direction of effect compared to the other areas of the cerebral cortex may indicate a reversed pathophysiological significance of the DLPFC compared to other cerebral cortical territories. For example, altered DLPFC-DN functional connectivity may represent a pathological abnormality in at-risk individuals who go on to develop psychosis, while the other observed alterations in functional connectivity may represent compensatory reorganizations, or vice versa. The same is true for the DN visual territory that revealed an opposite pattern of functional abnormality compared to the other two DN functional territories – abnormal functional connectivity in DN visual domains might contribute differently to the brain correlates of pre-clinical DN abnormal functional connectivity in schizophrenia when compared to DMN or salience-motor DN functional areas. In light of the recent reports on impairments in integration-segregation balance in schizophrenia (49), the results we report can be interpreted as DMN and motor parts of DN being more strongly integrated with the same systems in the cerebral cortex but more segregated with other systems (thus anticorrelated).

### Prior evidence linking psychosis risk and schizophrenia to cerebellum, as well as to DMN, salience-motor, and visual processing networks

Our study is the first to report abnormalities in functional connectivity of the DN within the cerebellum in this population, and also the first to report involvement of DMN, salience-motor, and visual networks in schizophrenia and psychosis risk by assessing these networks using seeds in the DN.

Cerebellar abnormalities are commonly reported in studies of schizophrenia. Structural cerebellar abnormalities in schizophrenia were identified decades ago (50), an observation that has been recently confirmed in a multi-site mega-analysis of close to one thousand patients (51). Moreover, PET (52) and fMRI (53) investigations detected abnormalities in cerebellar function, an observation that was later expanded to include non-schizophrenic subjects who are at an increased clinical (22, 23, 54, 55) or genetic risk (30, 56, 57) of developing schizophrenia. Cellular (34) and molecular (58) studies further support a role of the cerebellum in the pathophysiology of this disorder. Our study is the first to include the DN in the large and growing body of evidence linking cerebellum to schizophrenia, and the first to show DN abnormalities that precede the onset of cognitive and affective brain disease.

A large body of evidence has linked abnormalities in brain regions that belong to the DMN to schizophrenia, including studies that analyze resting-state functional connectivity (55, 59), task-activation (59–61), and structural data (see figure 4 in (51)). Other studies have also detected abnormalities not only within nodes of the DMN but also between DMN and other networks in schizophrenia, such as the salience network (62). Primary motor and sensory cortices have also been shown to exhibit abnormalities as indexed by functional (63) and structural analyses (64) (65) of patients diagnosed with schizophrenia or individuals at risk for developing this disease. Visual network abnormalities have also been detected in fMRI investigations of schizophrenia (66). A recent study using the same dataset reported abnormal functional connectivity in a broad range of functional territories in cerebral cortex preceding the onset of psychosis (38).

Our results provide further support for the notion that there is a wide range of functional networks implicated in the pathophysiology of schizophrenia, including in the mechanisms of disease that precede conversion to psychosis in individuals at risk, which can be captured by analyzing whole-brain functional connectivity from DN. A recently developed technique, *lesion network mapping*, assessed 89 brain lesions causing hallucinations and attributed DN for lesions causing auditory hallucinations (67). This finding may indicate that functional abnormalities of DN are especially relevant to the concurrent pathophysiology of emerging psychosis in individuals at risk. Recent developments in the field of neuromodulation and neurostimulation suggest that it may be possible in the near future to target specific sub-territories in human DN non-invasively (36, 68–71), hinting at potential future therapeutic interventions focused on DN sub-territories for the treatment or prevention of psychosis.

## Acknowledgements

This study was supported by the Ministry of Science and Technology of China (2016 YFC 1306803), US National Institute of Mental Health (R21 MH 093294, R01 MH 101052, and R01 MH 111448), and the European Union’s Horizon 2020 research and innovation program under the Marie Sklodowska-Curie grant agreement No. 749201 (to GC); MSK was supported by an NIMH Grant (R01 MH 64023); MES and RWM were supported by a VA Merit Award.

## Author contributions

LJS passed away on 7 September 2017 and RWM passed away on 27 May 2017. LJS and RWM were two of the initiators and principal investigators of the Shanghai At Risk for Psychosis (SHARP) study.

